# Mathematical Methods for Modeling Chemical Reaction Networks

**DOI:** 10.1101/070326

**Authors:** Justin Carden, Casian Pantea, Gheorge Craciun, Raghu Machiraju, Parag Mallick

## Abstract

Cancer’s cellular behavior is driven by alterations in the processes that cells use to sense and respond to diverse stimuli. Underlying these processes are a series of chemical processes (enzyme-substrate, protein-protein, etc.). Here we introduce a set of mathematical techniques for describing and characterizing these processes.

## I. Introduction to the type of problem in cancer

Cancer can be characterized by aberrations in regulatory mechanisms and signal transduction processes that alter how cells sense and respond to diverse stimuli. While a quantitative understanding of the mechanistic details of how these aberrations drive oncogenesis remains elusive, computational modeling and simulation can provide some unique insights and predictions into how regulatory and metabolic processes ultimately give rise to diverse phenotypes (e.g. metastasis). Models of these processes may ultimately play a role in both understanding them, and in understanding how to impact them (e.g. through therapeutic intervention). One promising modeling approach represents the diverse aspects of cellular control, ranging from protein-protein interactions to enzyme processing of metabolites as networks of coupled chemical reactions.

## II. Quick Guide to the Methods

### A. Differential equations for reaction networks

The dynamics of reaction networks are modeled by systems of ordinary differential equations (ODEs) tracking the time evolution of chemical concentrations for the species in the network. This allows the study of the properties of the dynamics (e.g. stability of steady states, existence of multiple steady states, etc.), using theoretical results of dynamical systems, or numerical simulations.

The most common way to construct ODEs from reaction networks is via the law of *mass-action kinetics*, under which the rate of a reaction is proportional to the concentration of each reactant^1^ (the proportionality factor is a positive number called the *rate constant*). Every reaction network has a corresponding mass-action ODE system. To illustrate, consider reaction network (1) of an overall reaction S1 + S2 → *P*, under a mechanism for enzyme catalysis corresponding to unordered substrate binding^2^ [1]:

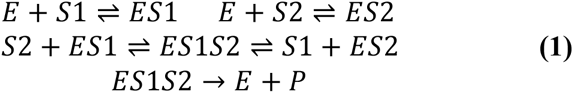

Network (1) includes seven species *E*, *S*1, *S*2, *ES*1, *ES*2, *ES*1*S*2, *P* and nine reactions among them (four reversible pairs and one irreversible reaction), whose rate constants are denoted by *k*_1_,…,*k_9_*. The corresponding mass-action system of ODEs is given in (2), and is obtained by collecting in the time derivative of each species the contribution of all reactions to the concentration of that species. For example, *E* is being produced in reaction *ES*1 → *E* + *S*1; this reaction has rate 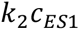 and we include this term in the expression of 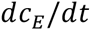 (the time derivative of concentration *c_E_* of *E*). On the other hand, *E* is consumed in the reaction *E* + *S*1 → *ES*1, therefore the rate 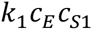 of this reaction is added with negative sign in the expression of 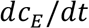. Note that reactions not involving *E* (e.g. *S*2 + *ES*1 → *ES*1*S*2) do not affect the concentration *c_E_*, and do not contribute terms to the expression of 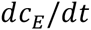. We assume additionally that substrates *S*1 and *S*2 are being supplied to the reaction environment at constant rates *F_S1_* and *F_S2_*, and we include these inflow (supply) rates into the expressions of 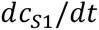 and 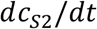 respectively. (In mass-action models, the inflow rates are always positive constants.) Moreover, the substrates *S*1, *S*2 and the product *P* are also being transported out of the reaction environment (or degraded) at rates 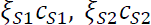 and 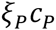 proportional to their concentrations (this is always the form of outflow rates in mass-action models, and the positive constants 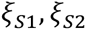 and ξ*_P_* are called outflow coefficients). The outflow terms are included with negative signs in the expressions of 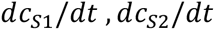 and 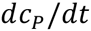 respectively.

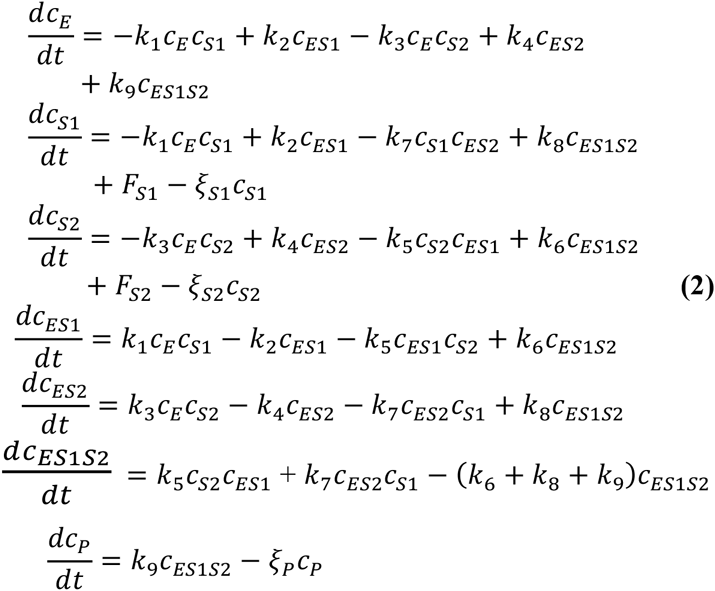

The differential equations (2) depend on 14 parameters (rate constants *k*_1_,…,*k*_9_ and the inflow/outflow rates 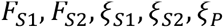). The presence of nonlinear terms in such a system warrants the possibility for complex behavior; indeed, it turns out that some choices of parameters^3^ give rise to *bistability*, i.e. there are two distinct steady states, both compatible with the same amount of enzyme (for example, such that *c_E_* + *c*_*ES*1_ + *c*_*ES*2_ + *c*_*ES*1*S*2_ = 1). For the particular choice of parameters, there are two stable steady states (depicted as red dots in Fig. 1) and one unstable steady state (green dot in Fig.1).

**Figure 1.**
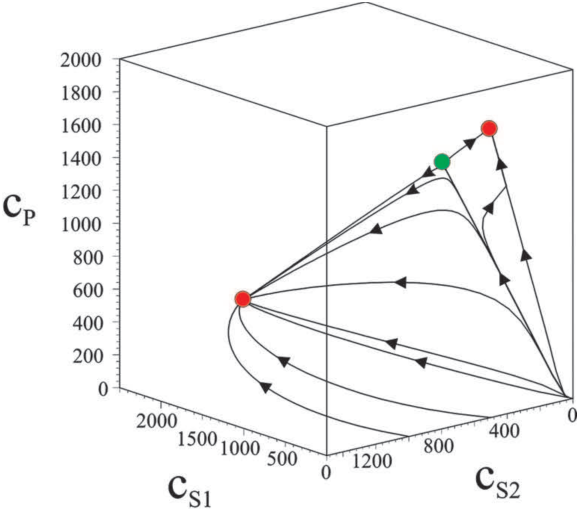
Examples of trajectories for S1, S2, P concentrations. Figure taken from [1].

The two stable steady states are reached from initial concentrations in their respective basins of attraction, and switching between them is possible, for example, by a temporary disturbance in a substrate supply rate.

Numerical simulation may reveal bistability of a network for given values of parameters (rate constants, inflow/outflow rates, enzyme total concentrations). In practice however, there are often no good estimates for parameter values, and sampling a vast parameter space in search of interesting behavior is almost an impossible task. However, in many cases, one can answer questions like “are there parameter values that give rise to bistability for a certain network?” without numerical simulation, but rather based on a body of theoretical work termed *reaction network theory*.

### B. Reaction network theory

Beginning with groundbreaking work of Horn, Jackson and Feinberg [3,4,5], reaction network theory addresses the question: “What behaviors of a reaction network are a function of its structure, and are robust to different choices of kinetics?” This question has a natural extension which is practically very important: “How can reaction networks be controlled in order to encourage/rule out certain behaviors?” The early work of Horn, Jackson and Feinberg solved a number of difficult problems on chemical reaction network behavior, for systems with mass-action kinetics.

Recently there has been an explosion of interest in reaction network theory, fueled by the growth of systems biology, and the development of new and powerful mathematical techniques. A very fruitful area of research has been in identifying conditions for multiple positive steady states in reaction networks [1,6,7,8,9]. In particular, the cycle structure of the *SR graph* [1] or *DSR graph* [7] associated to a network may rule out bistability for any choice of rate constants. The DSR graph is a directed labeled bipartite graph resembling the schematic of the reaction network. It turns out that the DSR graph of network (1), depicted in Fig.2, cannot rule out the possibility of multiple positive steady states. While this is not enough to insure that bistability is possible, further theory [6] shows that in fact there are choices of parameters for which bistability occurs.

**Figure 2.**
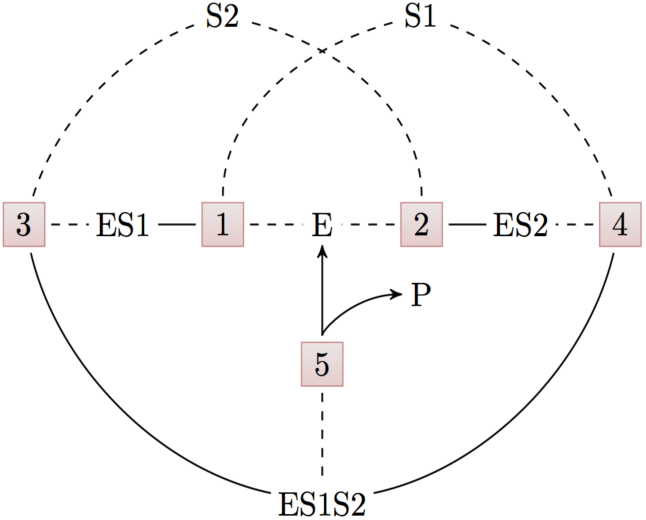
The DSR graph of network (1), generated using the software CoNtRol [10]

Methods like these show an intimate relationship between network structure and bistability, and provide some surprising and counterintuitive results. For example, Table 1 shows how bistability can be induced by subtle differences in enzyme catalysis mechanisms for the same overall reaction (for example, *S → P* in entries 3 and 4). The webserver CoNtRol [10] implements a series of theoretical methods [8] testing the possibility for bistability in reaction networks. Another powerful software is the Chemical Reaction Network Toolbox [11].

**TABLE 1.** Examples of enzymatic networks and their capacity for bistability. Table taken from [1].

| Entry | Network | Remark | Bistability |
| --- | --- | --- | --- |
| 1 | $E+S \rightleftharpoons ES \rightarrow E+P$ | Elementary enzyme catalysis underlying Michaelis-Menten kinetics: $S \rightarrow P$ | N |
| 2 | $E+S \rightleftharpoons ES \rightarrow E+P$, $E+I \rightleftharpoons EI$ | Elementary enzyme catalysis with competitive inhibition: $S \rightarrow P$ | N |
| 3 | $E+S \rightleftharpoons ES \rightarrow E+P$, $ES+I \rightleftharpoons ESI$ | Elementary enzyme catalysis with uncompetitive inhibition: $S \rightarrow P$ | N |
| 4 | $E+S \rightleftharpoons ES \rightarrow E+P$, $E+I \rightleftharpoons EI$, $ES+I \rightleftharpoons ESI \rightleftharpoons EI+S$ | Elementary enzyme catalysis with mixed inhibition: $S \rightarrow P$ | Y |
| 5 | $E+S1 \rightleftharpoons ES1$, $S2+ES1 \rightleftharpoons ES1S2 \rightarrow E+P$ | Two-substrate enzyme catalysis with ordered substrate-binding $S1+S2 \rightarrow P$ | Y |
| 6 | $E+S1 \rightleftharpoons ES1$, $E+S2 \rightleftharpoons ES2$, $S2+ES1 \rightleftharpoons ES1S2 \rightarrow S1+ES2$, $ES1S2 \rightarrow E+P$ | Two-substrate enzyme catalysis with un-ordered substrate-binding $S1+S2 \rightarrow P$ | Y |

<sup>3</sup> For example, $k_1 = 93.43$ , $k_2 = 2539$ , $k_3 = 481.6$ , $k_4 = 1183$ , $k_5 = 1556$ , $k_6 = 121192$ , $k_7 = 0.02213$ , $k_8 = 1689$ , $k_9 = 85842$ , $\xi_{S1} = \xi_{S2} = \xi_P = 1$ , $F_{S1} = 2500$ , $F_{S2} = 1500$ . The values are taken from [1], and they were used to draw Fig. 1.

Other questions in reaction network theory have also witnessed notable progress: *oscillations* [12,13]; *global stability and global convergence*, i.e., informally speaking, the question of when all initial conditions lead to the same stable behavior [14–18]; *persistence*, i.e., whether some reactant concentrations can decrease to zero despite positive initial concentrations [14,15,19,20].

## Footnotes

1 To be exact, proportional to a power of the reactant concentration given by its stoichiometry. For example, the rate of a dimerization 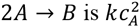.

2 This mechanism is involved in cyclin-dependent kinase-catalyzed phosphorylation reactions that play important roles in the cell cycle [2].

3 For example, *k*_1_ = 93.43, *k*_2_ = 2539, *k*_3_ = 481.6, *k*_4_ = 1183, *k*_5_ = 1556, *k*_6_ = 121192, *k*_7_ = 0.02213, *k*_8_ =1689, *k*_9_ = 85842, *ξ*_*S*1_ = *ξ*_*S*2_ = *ξ_p_* = 1, *F*_*S*1_ = 2500, *F*_*S*2_ = 1500. The values are taken from [1], and they were used to draw Fig. 1.

